# ACE: A Workbench using Evolutionary Genetic Algorithms for analysing association in TCGA Data

**DOI:** 10.1101/274175

**Authors:** Alan Gilmore, Kienan I Savage, Paul O’Reilly, Aideen C Roddy, Philip D Dunne, Mark Lawler, Simon S McDade, David J Waugh, Darragh G McArt

**Affiliations:** Centre for Cancer Research and Cell Biology, Queen’s University Belfast, 97 Lisburn Road, Belfast, Northern Ireland, BT9 7AE, United Kingdom

**Keywords:** Evolutionary algorithm, visualisation, biomarker correlation

## Abstract

Modern methods in generating molecular data have dramatically scaled in recent years, allowing researchers to efficiently acquire large volumes of information. However, this has increased the challenge of recognising interesting patterns within the data. *Atlas Correlation Explorer* (*ACE*) is a user-friendly workbench for seeking associations between attributes in the cancer genome atlas (*TCGA*) database. It allows any combination of clinical and genomic data streams to be selected for searching, and highlights significant correlations within the chosen data. It is based on an evolutionary algorithm which is capable of producing results for very large searches in a short time.

## Introduction

The capacity to sequence and profile multiple tumour/tissue samples rapidly and at affordable costs is making it easy for researchers to acquire large volumes of information. In turn, analysing or searching these collections in conventional ways can be exhaustive, and researchers often restrict their focus to areas already known from prior analyses, making it inevitable that many anomalies and insights hidden in this data will be overlooked (Wang et al. 2016).

In efforts to address these challenges we designed a software environment to enhance the ability of researchers to efficiently and safely explore such data collections for new biological insights. The framework employs sophisticated engineering methods which implementing evolution-based algorithms to improve the efficiency of data-mining. The initial implementation is based on *TCGA* (“The Cancer Genome Atlas”) (TCGA Research Network et al. 2013) data and was therefore coined *ACE* (“Atlas Correlation Explorer”).

The core technology of *ACE* is an engine which uses a genetic approach modelled on evolution to carry out its search. Genetic algorithms represent an exciting area of evolution-based logic, which seek to find near-optimal solutions using minimal resources in computationally-intensive problems and they have been applied in many differing fields (Notredame 1997, Aerts 2004, Tumuluru and McCulloch 2016).This engine creates a large pool of software “organisms”, each of which looks for correlation between features in *TCGA* data. As with real-world evolution it is indiscriminate in the initial selection, so in the vast majority of cases no correlation is found. However, when any correlation is found, that organism will thrive in the evolutionary pool, and will be refined through the equivalent of mating and mutation. The engine creates vast numbers of such organisms, and once evolution has been allowed to continue for some time, the best organisms will have identified some significant connections. This engine was originally developed to address pattern analysis applications in financial markets, where the problems were too complex to be addressed by “brute force” methods, and a genetic approach was found to allow convergence to near-optimal solutions in practical timescales. The key strength is that the engine can uncover biological relationships in the given ‘population’ under study that a human expert would be unlikely to observe.

## ACE

In the case of ACE, some additional techniques have been used to enhance performance. The data used by the engine was derived from the Broad Firehose TCGA pipelines (Broad Institute TCGA Genome Data Analysis Center 2016), but has been reformatted into pre-processed binary files that can be loaded directly into memory for very fast access. Data associated with successful “organisms” are cached for re-use. ACE is implemented as a *Windows* desktop application with requirements of *Windows 7* or later (Figure 1). ACE was developed using C# and *Microsoft Visual Studio*. Approximately two gigabytes of space are required for the converted data of each cancer type. In using *ACE*, a researcher can select subsets of the *TCGA* data, and *ACE* will search the values in those subsets and pick out those that show strong correlation. When the chosen subsets are large, an exhaustive search for the best correlations would be impractical, but the evolutionary nature of *ACE* allows for good associations to be found in reasonably short times.

This engine is flexible in application, allowing systems using it to define their own “universe”, to define the type of organisms that will exist within it, and to define mutation and mating algorithms relevant to that universe. The “survival of the fittest” principle is used, but each system using the engine can define a scoring system for organisms, letting only the highest-scoring individuals survive to produce offspring. This allows a system implemented using the engine complete flexibility in setting goals for the evolution.

ACE has two main screen views, and a user can toggle between them:

- The Data Selection View allows browsing of the TCGA data, and selection of portions of it to be analysed by ACE.
- The Genetic Search View is used to run an evolution based on the selected data, and to view a ‘leaderboard’, showing the best correlations found by ACE between items of the selected data.

An import and conversion phase is needed to prepare the TCGA data for each cancer type for use with ACE. Currently the following cancer types are supported: Bladder urothelial carcinoma (BLCA); Breast invasive carcinoma (BRCA); Colon adenocarcinoma (COAD); Glioblastoma multiforme (GBM); Head and Neck squamous cell carcinoma (HNSC); Lung adenocarcinoma (LUAD); and Prostate adenocarcinoma (PRAD). Additional cancer types will be included in future releases and a comprehensive manual can be found in the supplementary materials and methods.

**Figure 1.**
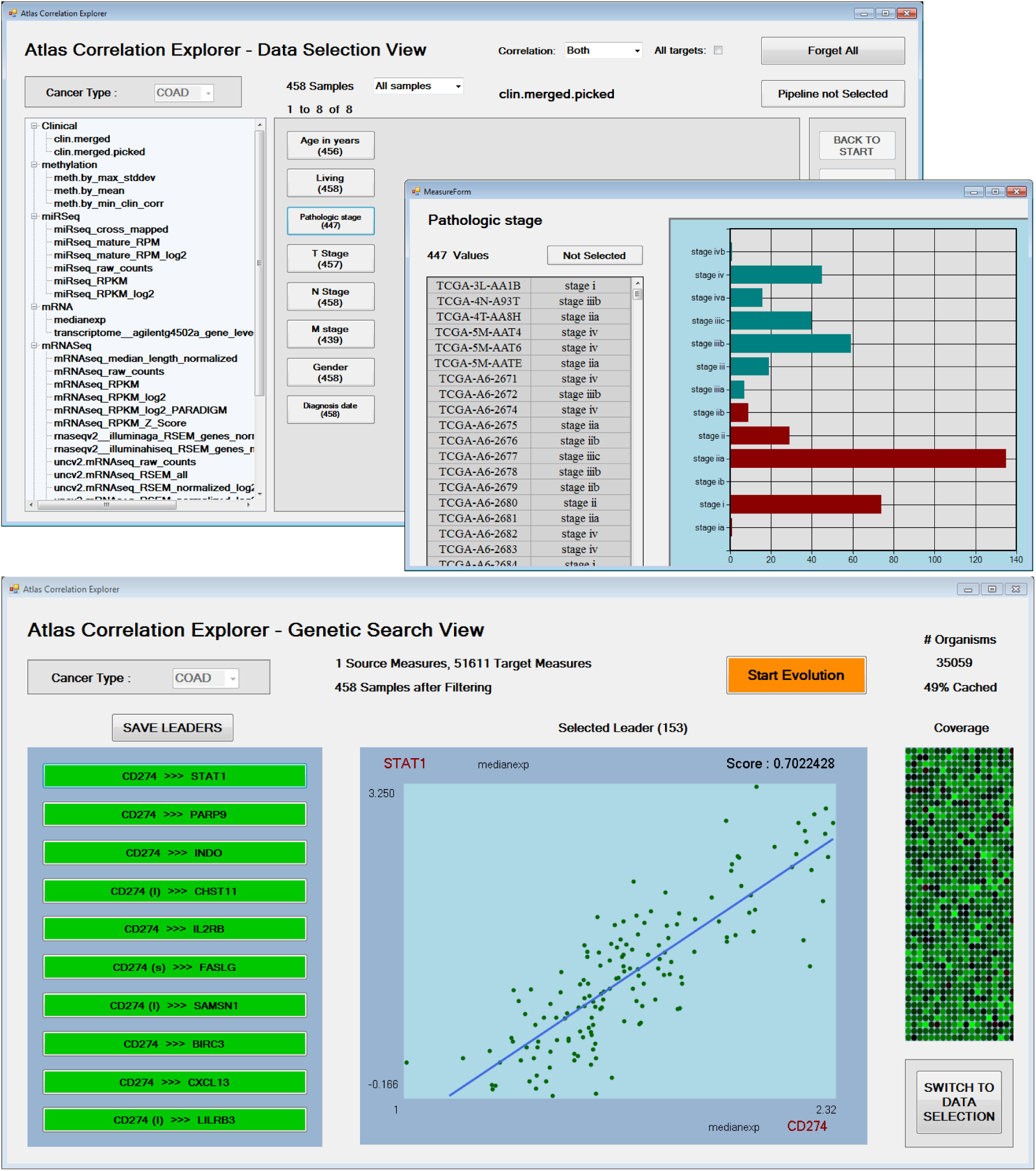
**Top:** This illustrates the ‘Data Selection View’. In this case the cancer type being studied is Colorectal Adenocarcinoma (COAD), and the area of interest highlighted for possible thresholding is the pathology stage. **Bottom:** This demonstrates the Genetic Search View being used to search for correlation between expression, methylation, protein and mutational data with CD274 transcription as the marker of interest. The list generated dynamically on the left hand side represents the best correlations found so far, and the main panel shows strong association depicting top hits, STAT1, PARP9 and INDO. The (s) indicates that a correlation has been found using the SQUARE ROOT of the given values. As part of the evolution, this is tried in addition to LOG (l) and ARCSIN (a) as investigation has shown that these sometimes provide a better fit than the raw values.

## Supplementary information

Supplementary data regarding installation and utility are available on GitHub along with the current software version.

## Availability and Implementation

ACE is freely available for non-commercial use as a binary download from: https://github.com/AlanRGilmore/ACE/releases

### Acknowledgments and funding information

This work was supported by a CRUK PhD studentship (A.R. and D.G.M.)

## References

Aerts S, Van Loo P, Moreau Y, De Moor B. 2004. A genetic algorithm for the detection of new cis-regulatory modules in sets of coregulated genes. Bioinformatics. 20 (12): 1974–1976.

Broad Institute TCGA Genome Data Analysis Center. 2016. Analysis-ready standardized TCGA data from Broad GDAC Firehose 2016_01_28 run. Broad Institute of MIT and Harvard. Dataset. https://doi.org/10.7908/C11G0KM9

Notredame C, O’Brien EA, Higgins DG. 1997. RAGA: RNA sequence alignment by genetic algorithm. Nucleic Acids Res. 25 (22): 4570–4580.

The Cancer Genome Atlas Research Network, Weinstein JN, Collisson EA, Mills GB, Mills Shaw KR, Ozenberger BA, Ellrott K, Shmulevich I, Sander C, Stuart JM. 2013. The Cancer Genome Atlas Pan-Cancer analysis project. NatGenetics.45: 1113–1120.

Tumuluru JS, McCulloch R. 2016. Application of Hybrid Genetic Algorithm Routine in Optimizing Food and Bioengineering Processes. Foods. 5(4): 76.

Wang L, Wang Y, Chang Q. 2016. Feature selection methods for big data bioinformatics: A survey from the search perspective. Methods. 111: 21–31

